# Swallowing-related neural oscillation: An intracranial EEG study

**DOI:** 10.1101/2020.12.05.412783

**Authors:** Hiroaki Hashimoto, Kazutaka Takahashi, Seiji Kameda, Fumiaki Yoshida, Hitoshi Maezawa, Satoru Oshino, Naoki Tani, Hui Ming Khoo, Takufumi Yanagisawa, Toshiki Yoshimine, Haruhiko Kishima, Masayuki Hirata

## Abstract

Swallowing is a unique movement due to the indispensable orchestration of voluntary and involuntary movement. The transition from voluntary to involuntary swallowing is executed on the order of milliseconds. We hypothesized that its neural mechanism is revealed by high frequency cortical activities. Eight epileptic participants fitted with intracranial electrodes over the orofacial cortex were asked to swallow a water bolus, and cortical oscillatory changes, including high γ band (75–150 Hz) and β band (13–30 Hz) were investigated at the time of mouth-opening, water-injection, and swallowing. High γ power increases associated with mouth-opening were observed in the ventrolateral prefrontal cortex with water-injection in the lateral central sulcus and with swallowing in the region along the Sylvian fissure. Mouth-opening induced a β power decrease, which continued until the completion of swallowing. The high γ burst activity was focal and specific to swallowing, however, the β activities were extensive and not specific to swallowing. At the boundary time between voluntary and involuntary swallowing, swallowing-related high γ power achieved the peak, and subsequently, the power decreased. We demonstrated three distinct activities related to mouth-opening, water-injection, and swallowing induced at different timings, using high γ activities. The peak of high γ power related to swallowing suggests that during voluntary swallowing phases, the cortex is the main driving force for swallowing rather than the brain stem.

## 1. Introduction

Swallowing is a primitive and fundamental function, and also a unique movement because the cooperation between voluntary and involuntary movements are necessary for normal swallowing. Thus, the swallowing movement is divided into two components, voluntary movement and involuntary movement (Jean, 2001). The cerebral cortex is assumed to play a crucial role in swallowing movements along with the brain stem. The cerebral cortex triggers swallowing and modulates the brain stem where the central pattern generator (CPG) for swallowing is assumed to be in humans (Ertekin and Aydogdu, 2003). Therefore, damage of the cerebral cortex due to stroke (Marik, 2001; Smithard et al., 2007) or neurodegenerative disease (Boccardi et al., 2016; Jani and Gore, 2016) promotes swallowing disturbance (dysphagia).

In clinical practice as treatments for dysphagia, adopting different postures and modifying the bolus consistency are widely accepted (Speyer et al., 2010). Peripheral electrical stimulation, such as pharyngeal electrical stimulation (Suntrup et al., 2015) or transcutaneous neuromuscular stimulation (Ludlow et al., 2007), and non-invasive brain stimulation using repetitive transcranial magnetic stimulation (rTMS) (Verin and Leroi, 2009) or transcranial direct current stimulation (tDCS) (Vasant et al., 2014) are also being studied as emerging therapeutic strategies. In these neuromodulation strategies for dysphasia, positive effects have been reported, however, various intensities and durations were used for stimulation (Sasegbon et al., 2020), and the best condition for stimulation remains unclear. Therefore, the treatment effects of these neuromodulation therapy remain limited, and further studies are required to provide evidence for overcoming dysphagia.

We have continued to research in order to contribute to the field of neuromodulation for dysphagia, using our brain-machine interface (BMI) technology. First, we newly developed a swallow-tracking system that enabled us to monitor swallowing noninvasively and simultaneously, using intracranial electroencephalograms (EEGs) (Hashimoto et al., 2018). Next, we demonstrated that deep transfer learning that were applied to intracranial EEG signals well classified swallowing intention (Hashimoto et al.). Current neuromodulation strategies for dysphagia stimulate brain areas or peripheral organs regardless of patients’ swallowing intention. We guess that if the stimulation is executed by patients’ swallowing intention, more recovery will be obtained by sensory neurofeedback. BMI technology is useful for the realization of stimulation triggered by patients’ intention.

The elucidation of swallowing-related neural processing is essential for the realization of such a swallow-assisting BMI. Non-invasive methods, such as scalp EEG, (Jestrovic et al., 2017; Yang et al., 2014) positron emission tomography (PET), (Hamdy et al., 1999b) near-infrared spectroscopy (NIRS) (Kober and Wood, 2014), TMS, (Hamdy et al., 1996) functional magnetic resonance imaging (fMRI), (Hamdy et al., 1999a; Martin et al., 2001; Toogood et al., 2017a) and magnetoencephalography (MEG) (Dziewas et al., 2003; Furlong et al., 2004) implicate multiple cortical sites involved in swallowing. fMRI has high spatial resolution on the order of millimetres; however, it has poor temporal resolution (Aine, 1995). The temporal resolution of MEG is high, on the order of milliseconds (Furlong et al., 2004); however, its spatial resolution is limited. Non-invasive methods have either weak temporal or spatial resolution, therefore, we inferred that such methods had a potential limitation for investigation of neural processing related to swallowing that transits from voluntary to involuntary phase on the order of milliseconds (Ertekin, 2002; Ertekin and Aydogdu, 2003; Nascimento et al., 2015).

Intracranial EEG, such as electrocorticogram (ECoG), that record neural activities directly from the cortical surface of the brain enable us to measure neural activities with high spatiotemporal resolution. Moreover, using ECoG, we can measure high frequency neural oscillations, including high γ band (> 50 Hz). High γ activity is a key oscillation that reflects the neural processing of sensory, motor, and cognitive events (Canolty et al., 2006; Crone et al., 1998; Hashimoto et al., 2017). High γ activities also show better functional localization than lower-frequency bands (Crone et al., 1998), and have come to be known as a useful features for decoding of neural signals (Hashimoto et al.). Here, we hypothesized that analysis of high γ activities acquired by ECoG measurement would be able to reveal the swallowing-related cerebral oscillation changes on the order of milliseconds that remained unclear. Novel findings may provide neuromodulation strategies for dysphagia with beneficial information, such as brain areas and timing for stimulation that enable us to achieve better results.

## 2. Methods

### 2.1. Participants

Eight patients with intractable temporal lobe epilepsy participated in this study (four females, 10’s–50’s years of age) (Table 1). They were admitted to Osaka University Hospital from April 2015 to July 2019, and underwent intracranial electrodes placement for invasive EEG study. They had no swallowing disturbances, which was confirmed by medical interview. All participants or their guardians were informed of the purpose and possible consequences of this study, and written informed consent was obtained. The present study was approved by the Ethics Committee of Osaka University Hospital (Nos. 08061, 16469).

**Table 1.**
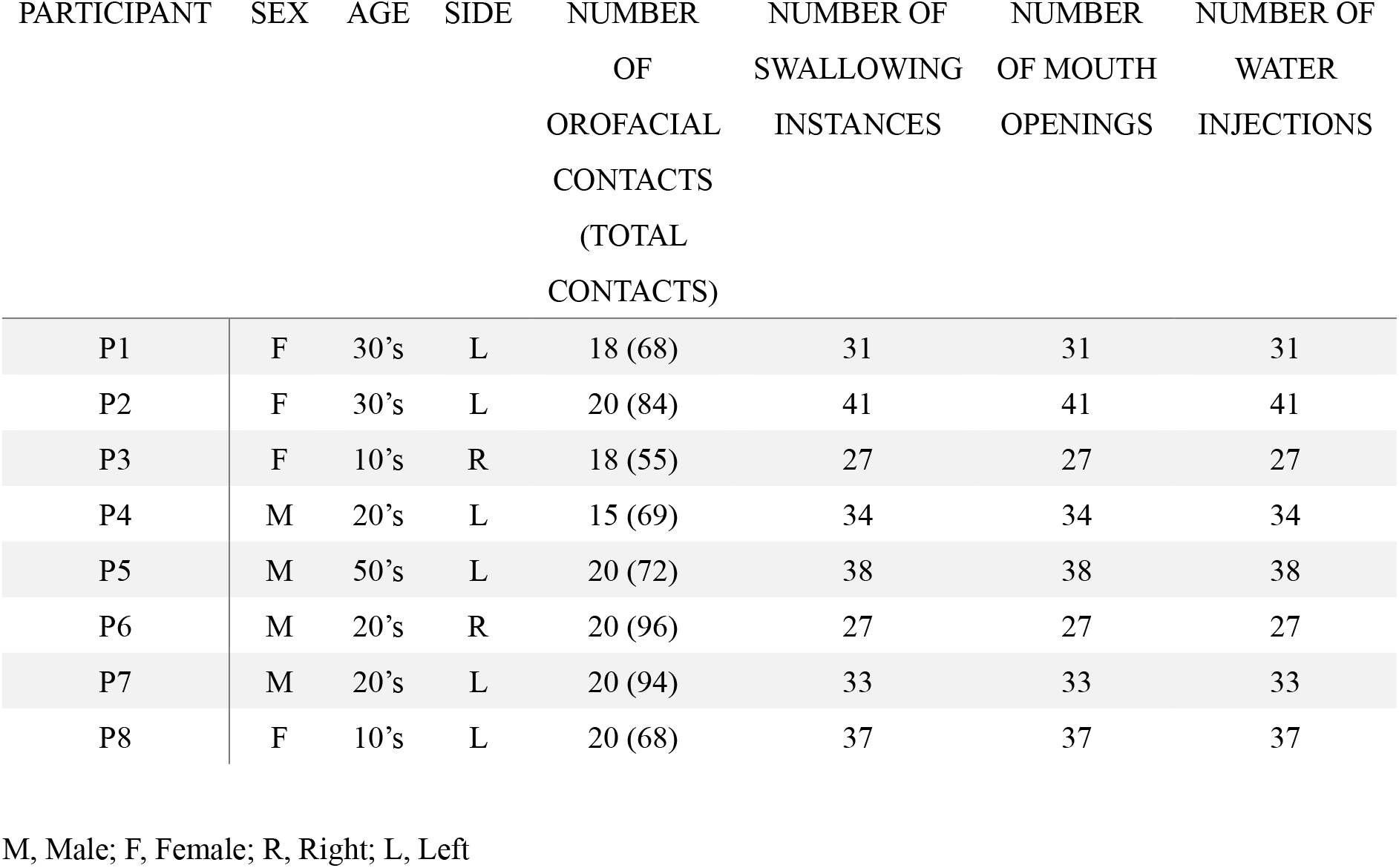
Clinical profiles

### 2.2. Intracranial electrodes

Different types of electrodes (Unique Medical Co. Ltd., Tokyo, Japan), including grid, strip, and depth types, were implanted in the subdural space during conventional craniotomy as a clinical epileptic surgery. For analysis, we chose planar-surface platinum grid electrodes (4×5 contacts array) that were placed overt the lateral portion of the central sulcus corresponding to the orofacial cortex. The number of total implanted contacts and selected contacts in each participant are shown in Table 1. The diameter of the contacts was 3 or 5 mm, and the intercontact center-to-center distance was 5, 7, or 10 mm.

### 2.3. Electrode location

Preoperative structural MRI was obtained with a 1.5-T or 3.0-T MRI scanner, and postoperative computed tomography (CT) scans were acquired with the implanted electrodes in place. The implanted electrodes that were obtained from the CT images were overlaid onto the 3-dimensional brain renderings from the MRI volume that was created by FreeSurfer software (https://surfer.nmr.mgh.harvard.edu). We obtained the Montreal Neurological Institute (MNI) coordinates of the implanted contacts with Brainstorm software (http://neuroimage.usc.edu/brainstorm/), which merged the preoperative MRI scans and postoperative CT scans. The location of the implanted electrodes that was visualized by Brainstorm was then confirmed by intraoperative photographs.

### 2.4. Task

The experiments were performed approximately 1 week after surgical electrode placement when all participants fully recovered from surgery. The participants were asked to sit on a chair and to remain still, especially without moving their mouth. The participants then opened their mouths, and the examiner injected 2 mL of water into their mouths with a syringe. We requested that the participants keep the water bolus in their mouths for approximately 1 s and then swallow it at their own pace without external cueing to prevent erroneous volitional water swallowing (aspiration). After we confirmed that the participants had completed one swallowing movement, the next water bolus was injected.

### 2.5. Swallowing monitoring

For noninvasive swallowing detection, we used an electroglottograph (EGG), a microphone, and a motion-tracking system (Fig.1A). A laryngograph (Laryngograph Ltd, London, UK) was used as an EGG and recorded the neck impedance changes associated with swallowing (Firmin et al., 1997) (Fig.1B). A pair of contacts was placed on the neck skin below the thyroid cartilage of the participants at an intercontact center-to-center distance of 25 mm and was held in place by an elastic band.

**Fig 1.**
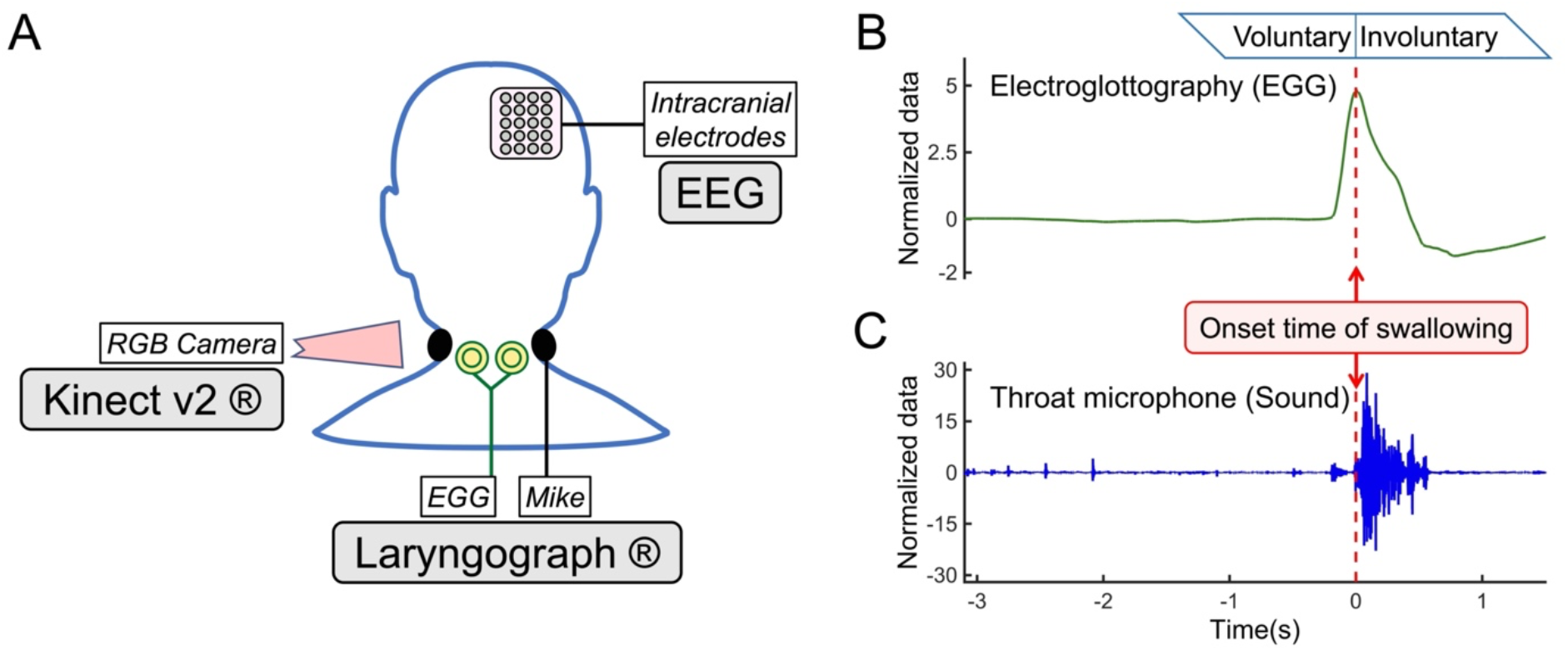
Multimodal data related to swallowing. Intracranial EEG were recorded during participants swallowed. The swallowing was monitored by an RGB camera of Kinect v2, and an electroglottograph (EGG) and a microphone of Laryngograph (A). Across-trials averaged impedance waveforms of an EGG (B) and a throat microphone (sound) (C), from one participant (P1) are shown. For analysis, the onset of swallowing was defined as the peak time of an impedance waveform. The onset time corresponds to the boundary time between voluntary and involuntary swallowing.

Sounds of swallowing due to the bolus passing through the pharynx were detected by a throat microphone (Kusuhara et al., 2004) (Fig.1C). We connected the throat microphone (Inkou mike; SH-12iK, NANZU, Shizuoka, Japan) to the laryngograph to record the swallowing sounds. The shape of the microphone was arched to fit around the participant’s neck. The sampling rate of a laryngograph and a throat microphone was 24,000 Hz.

We captured the motion of the participants at 30 frames per second with the motion-tracking system, which was newly developed by us using Kinect v2 (Microsoft, Redmond, Washington, USA) (Hashimoto et al., 2018). The participants were seated facing Kinect v2, which was placed on a tripod at a distance of one meter, and their mouth and throat movements were captured automatically.

An electric stimulator (NS-101; Unique Medical, Tokyo, Japan) supplied digital synchronizing signals to the laryngograph and a 128-channel digital EEG system. The signals made an LED light flash, which was captured by the RGB camera of Kinect v2. The digital triggers and LED light flash enabled us to synchronize the multimodal data of an EGG, a microphone, the video captured by the RGB camera, and an intracranial EEG. The multimodal data enabled us to noninvasively monitor the swallowing movements. The video captured by the RGB camera enabled us to the detect time when the participants opened their mouths and when the water bolus was injected into the mouth.

### 2.6. Signal segmentation based on swallowing-related events

The swallowing onset time was determined at the time when the impedance waveform reached the peak (Fig.1B). Swallowing sounds occurred frequently as the bolus of water passed through the pharynx (Kusuhara et al., 2004), and their evaluation in conjunction with the EGG helped us to judge whether the impedance change was caused by swallowing (Fig.1C). Additionally, we confirmed that the changes in impedance and sounds corresponded to water swallowing by using the video captured by the RGB camera. We inserted swallowing triggers, which corresponded to the swallowing onset time, into the ECoG data.

A previous study associated the neck impedance changes with the following swallowing stages: stage I (the oral stage), stage II (the pharyngeal stage), and stage III (the esophageal stage). The time of the EGG peak corresponded to the beginning of the pharyngeal stage (Kusuhara et al., 2004). Since the oral phase is voluntarily controlled and the pharyngeal and esophageal phases are involuntarily controlled (Jean, 2001), the swallowing onset time in this study corresponded to the transition time from voluntary swallowing to involuntary swallowing (Fig.1B).

We could also detect the time when the participant opened his/her mouth and when the examiner injected water into the participants’ mouth using the video captured by the RGB camera. Mouth triggers and water triggers, which corresponded to different times, were also inserted into ECoG data (the number of each trigger is shown in Table 1).

### 2.7. Data acquisition and preprocessing

The ECoG signals were measured with a 128-channel digital EEG system (EEG 2000; Nihon Kohden Corporation, Tokyo, Japan) and digitized at a sampling rate of 1000 Hz with a 0.3 to 333 Hz bandpass filter to prevent aliasing, and a 60-Hz notch filter to eliminate the AC line noise. Before any further processing, contacts containing external noise or epileptic discharge were excluded from further analyses. The ECoG signals were digitally re-referenced to a common average of implanted total contacts in each participant. We analyzed the ECoG signals that were time-locked to the triggers.

Throughout the following analyses, a bandpass filter using a two-way least-squares finite impulse response filter (pop_eegfiltnew.m from the EEGLAB version 14.1.2b, https://sccn.ucsd.edu/eeglab/index.php) was applied to the ECoG signals. To prevent edge-effect artifacts, additional 250 msec data remained at the initial and end points of the pre band-pass filtered signals. After band-passed filtering, the 250 msec extra data were excluded.

### 2.8. Spectral analyses

A time-frequency analysis of the time-locked ECoG signals to each trigger was performed using EEGLAB with a frequency range from 1–330 Hz and spectral power (in dB) calculated in 0.5-Hz bins with 200 data-points (from −5.0 to 2.5 s in every 33-msec window). The baseline for the time-frequency analysis was initially 0.5 s.

To create power contour maps, the power of each contact was constructed from the preprocessed ECoG signal by using a band-pass filter including high γ band (75–150 Hz) and β band (13–30 Hz) in combination with the Hilbert transformation (Cohen, 2008). We calculated the averaged power during 0.5 s time-window in these two bands, and the averaged power was normalized with the mean and standard deviation of the power during −5.0 to −4.5 s of the swallowing triggers in Figure 2 and during −1.0 to −0.5 s of the mouth triggers in Figure 3.

**Fig 2.**
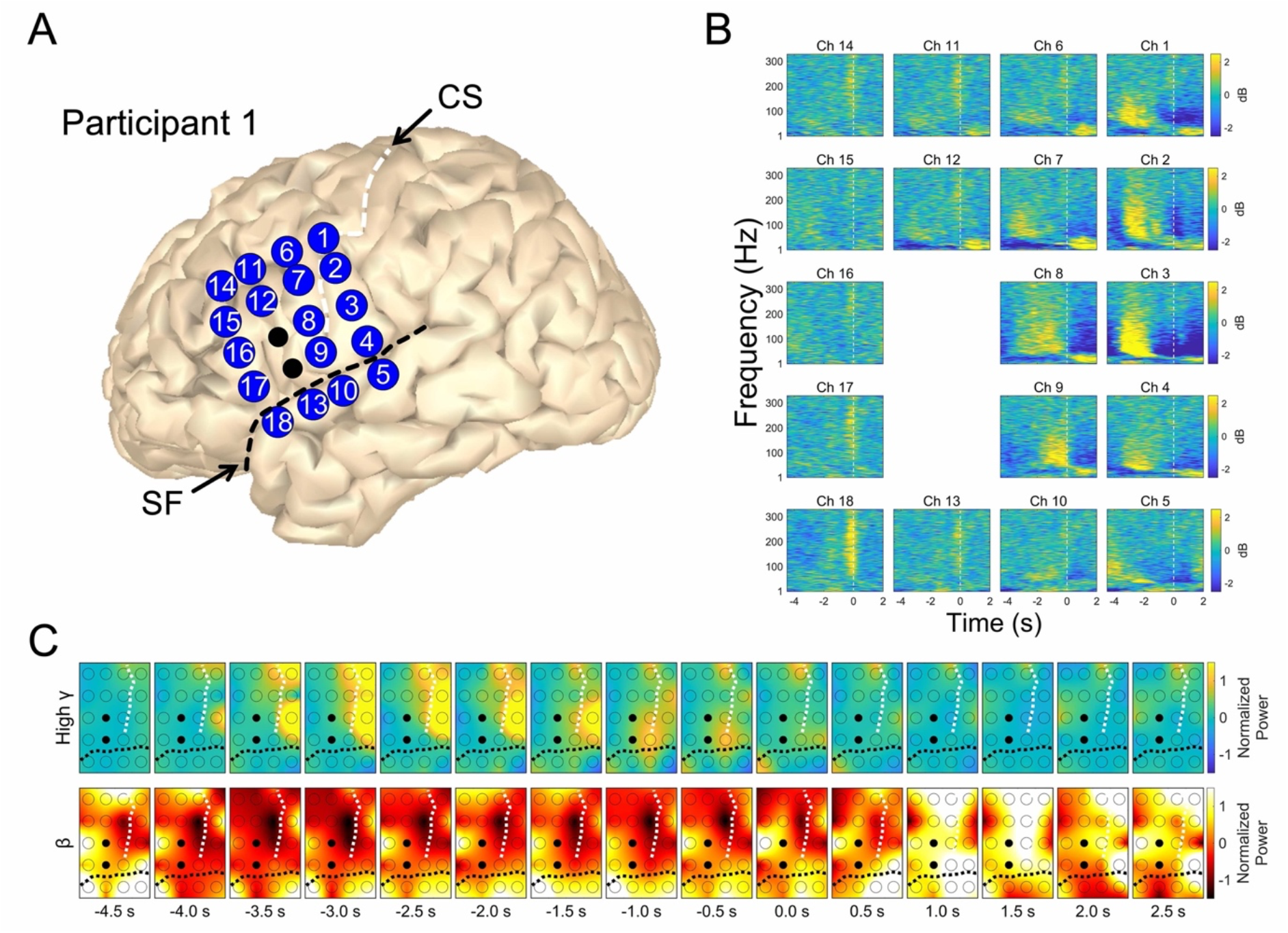
Temporal profiles of oscillatory changes associated with swallowing in P1. (A) The reconstructed MR images. The numbers correspond to the contact numbers. (B) Averaged time-frequency maps are shown from −4.5 to 2.0 s around the swallowing trigger. These baselines of 0.5 s duration range from −5.0 to −4.5 s. High γ band power (>50 Hz) increases specific to swallowing appeared in Ch 9 attached to the subcentral area from −2.0 to 0 s. Within −3.5 to −2.0 s, the time-frequency plots from the pre and postcentral gyri showed high γ increases along with decreases in lower frequency band (< 50 Hz) power (Ch 1, 2, 3, 4, 7, and 8). After 0 s, the pre- and postcentral gyri showed the lower frequency power increase (Ch 1, 2, 3, 4, 6, 7, 8, 9, and 12). (C) Contour maps of normalized power in high γ (75–150 Hz) and β (13–30 Hz) bands are shown from −4.5 to 2.5 s every 0.5 s interval around the swallowing trigger. The cortical areas in which the large power increased in the high γ band showed sequential changes. The contact that showed the largest high γ power increases immediately before swallowing onset time (−0.5 s) was Ch9, attached to the subcentral area. The two black contacts were excluded because of severe noise contamination. The central sulcus (CS) and the Sylvian fissure (SF) are indicated by the white and black dashed lines in each.

**Fig 3.**
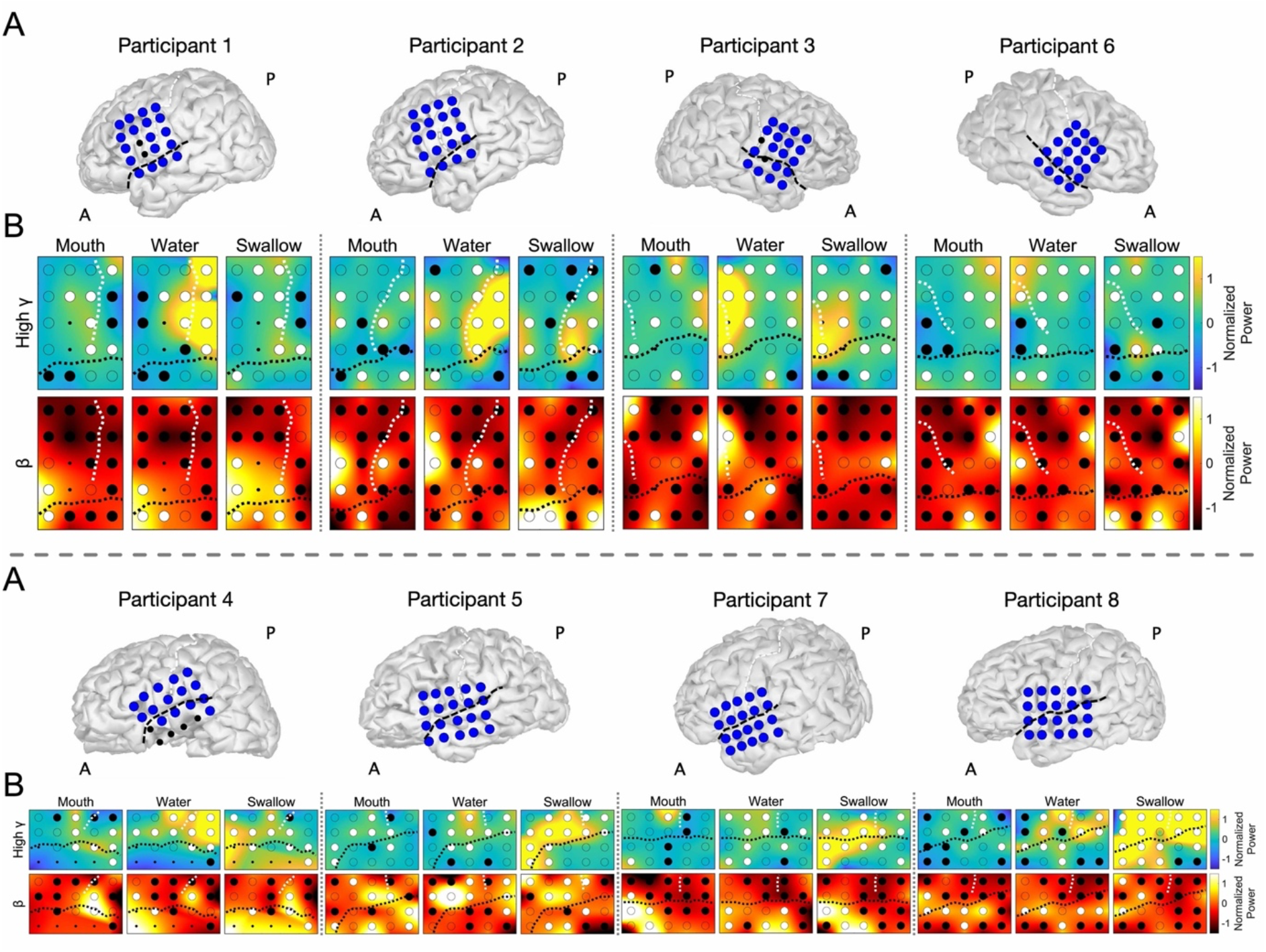
Power contour maps at mouth, water, and swallowing triggers in high γ and β band. (A) The reconstructed MR images for all participants. (B) The upper columns indicate high γ contour maps and the lower columns indicate β contour maps. Significant power increasing was indicated as white filled circles, and significant power decreasing was indicated as black filled circles (single-sided permutation test with Bonferroni correction, corrected *p* < 0.05). Excluded contacts were indicated as small black filled circles. The central sulcus and the Sylvian fissure are indicated by white and black dashed lines, respectively. Within the mouth, water, and swallowing, the mainly region where high γ burst was observed were the precentral gyrus, the postcentral gyrus, and the cortex along the Sylvian fissure in each. The β attenuation were observed in wide area related to all three triggers.

### 2.9 Extraction of contacts

Significant power increasing or decreasing contacts (single-sided permutation tests with Bonferroni correction) are indicated as filled white or black circles in Figure.3B. These contacts were plotted over the left hemisphere of MNI normalized brain using Brainstorm software (Fig.4A). The contacts attached to the right hemisphere were transposed to the left hemisphere.

**Fig 4.**
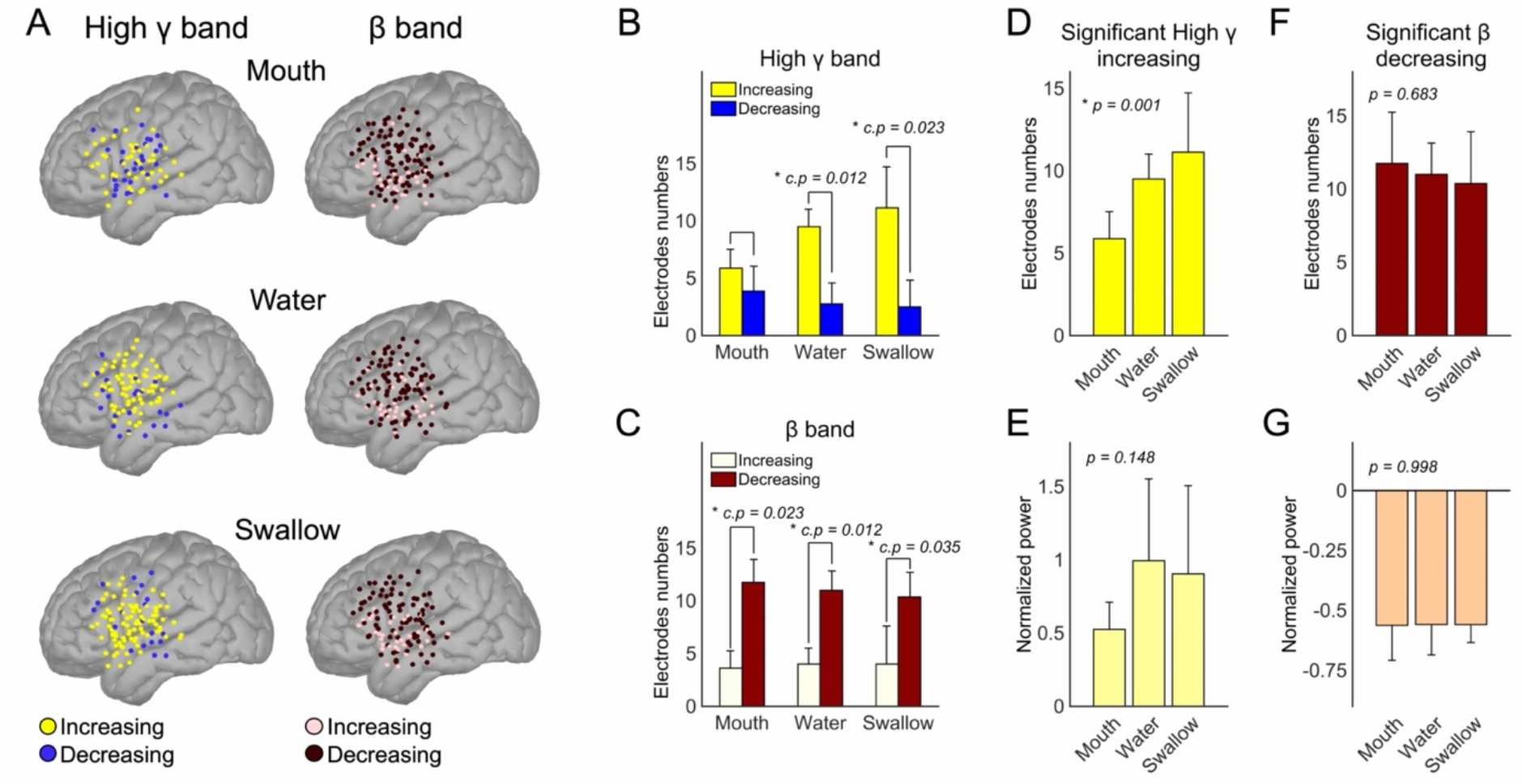
Profiles of high γ and β band related to mouth, water, and swallowing triggers. For each trigger group, contacts indicating significant power increases or decreases in high γ or β band were plotted over the left hemisphere of the MNI brain. In the high γ band, a power increase was notable, and in the β band, power decreasing was notable and extensive (A). Within the mouth, water, and swallowing group, the numbers of contacts were compared between the two groups including power increasing or power decreasing (single-sided Wilcoxon signed-rank test). P values were corrected with Bonferroni correction and significant p values (< 0.05) are shown indicated by an asterisk (corrected p: c.p). In the high γ band, the numbers of increasing contacts were significantly larger than that of decreasing contacts in the water and swallowing groups (B). In the β band, the numbers of decreasing contacts were significantly larger in all groups (C). In high γ power increasing contacts or β power decreasing contacts, the numbers of contacts and the degree of normalized power changing were compared across the mouth, water, and swallowing group (one-way ANOVA, significant *p* values (< 0.05) are indicated by with an asterisk) (high γ band in D and E, β band in F and G; the numbers of contacts in D and F, the degree of power changing in E and G). In the numbers of contacts with high γ increases, there were significant differences. There were no statistical differences in the normalized power changes in high γ band (E) and β band (G), and in the numbers of contacts in β band (F). The error bars indicate standard deviation (B-G).

The top 25% of contacts indicating a significant high γ power increase or a significant β power decrease were extracted and plotted on the left hemisphere of MNI normalized brain (Fig.5). These group of contacts associated with each trigger were defined as mouth-related contacts (Mouth-C), water-related contacts (Water-C), and swallowing-related contacts (Swallow-C).

**Fig 5.**
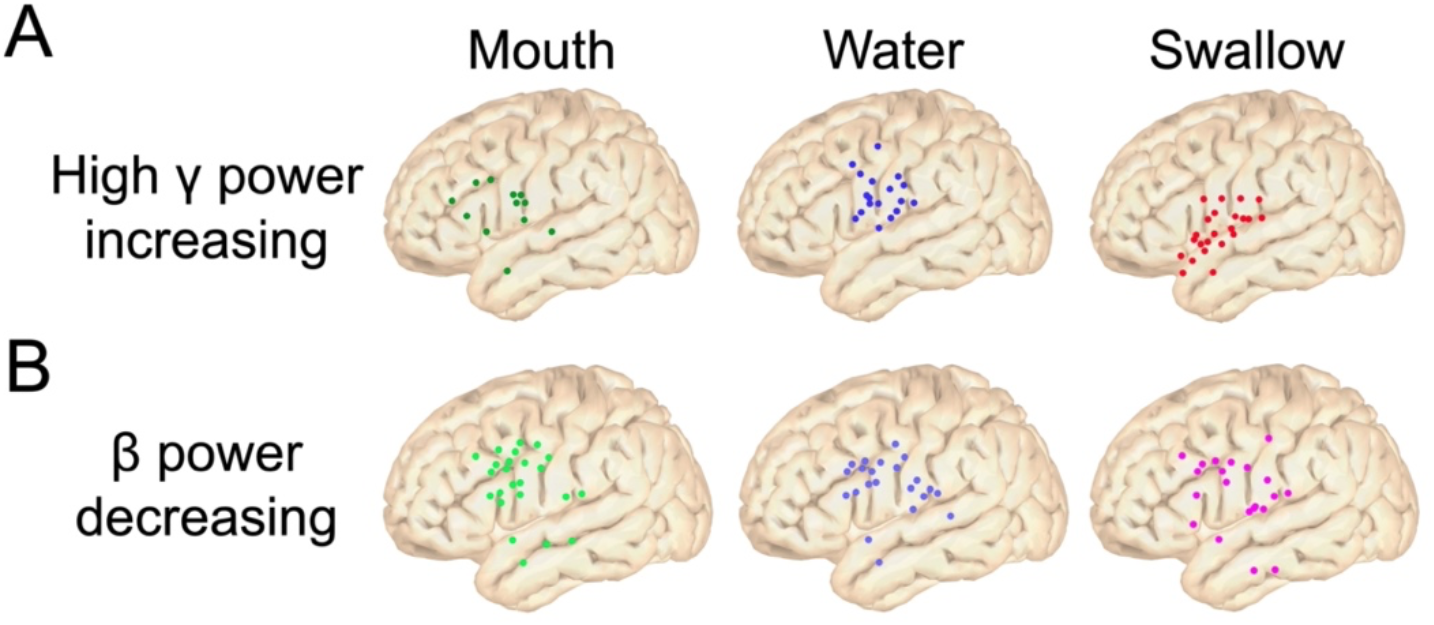
Spatial profile related to mouth, water, and swallowing trigger groups. The top 25% contacts indicating significant high γ power increases or β decreases were plotted over the left hemisphere of the MNI brain. In high γ increases, mouth-related contacts were observed in the precentral gyrus and the ventrolateral prefrontal cortex (VLPFC). Water-related contacts were observed in the lateral portion of the central sulcus, and swallowingrelated contacts were observed on the regions along the Sylvian fissure. In β decreasing, mouth- and water-related contacts were mainly localized in the VLPFC, and the localization loosen in the swallowing group.

### 2.10. Dynamic frequency power changes

We obtained averaged power waveforms of Mouth-C, Water-C, and Swallow-C relative to each trigger (mouth triggers, Mouth T; water triggers, Water T; swallowing triggers, Swallow T) in high γ and β band. A power time series were constructed from a band-pass filtered signal in combination with the Hilbert transformation (Cohen, 2008). The power time series were normalized by the power from base-time (−1.0 s to −0.9 s of Mouth T). Averaged frequency power values were calculated from a 100 msec time-window which was shifted every 10 msec. The shifted 100 msec data-points were compared with base-time data using the permutation test. Multiple comparisons were resolved by a family-wise error (FWE)-corrected threshold (see below).

### 2.11. Correlation analysis

We calculated Pearson correlation coefficients between normalized β power at 0 s of mouth triggers and sequential normalized β power from −1.5 s to 6 s of mouth triggers. Correlation coefficients were calculated in each participant, and then averaged. Using Monte Carlo method, we set the threshold of correlation coefficients that achieved 80% statistical power.

### 2.12. Statistics

For statistical evaluation of oscillatory power changes, we used a permutation test (Maris and Oostenveld, 2007). For comparison of two groups, single-sided Wilcoxon signed-rank test were used. For correction of multiple comparisons, we used Bonferroni correction or FWE-corrected threshold. For comparison of three groups of mouth triggers, water triggers, and swallowing triggers, we used one-way analysis of variance (ANOVA).

If one permutation test was done, we could obtain one distribution of differences. The maximum or minimum values of the distribution were stored, and the values at 99% of these distributions were taken as the FWE-corrected threshold, which is applied to the observed power time series (Cohen, 2014).

### 2.13. Data availability

All data that were generated by or analyzed in this study are available from the corresponding authors upon reasonable request and after additional ethics approvals regarding the data provision to individual institutions.

## 3 Results

### 3.1. Representative spatiotemporal oscillatory changes

The spatiotemporal oscillatory changes in a representative participant (Participant 1; P1) are shown in Figure 2. Averaged time-frequency maps around swallowing triggers from −4.5 to 2.0 s showed a high γ band (>50 Hz) power increase within about −3.5 to −2.0 s in the pre- and postcentral gyri (Fig.2A and 2B). Simultaneously, the pre- and postcentral gyri also showed a decrease in the low-frequency band <50 Hz. Within −2.0 to 0 s, a new high γ power increase was observed in Channel (Ch) 9. After 0 s, the pre- and postcentral gyri showed power increase in the low-frequency band <50 Hz.

A series of normalized power contour maps for the high γ band (75–150 Hz) and β band (13–30 Hz) were calculated from −4.5 to 2.5 s around swallowing triggers (Fig.2C). The location where the large high γ power increase appeared produced sequential changes. First, the large high γ power increase was observed in the pericentral gyri, especially for the postcentral gyrus during −4.0 to −1.5 s. Subsequently, the subcentral area, that is the narrow gyrus between the caudolateral extreme of the central sulcus and the Sylvian fissure, (Ch9) was the site where the large high γ power appeared during −1.0 to −0.5 s. Conversely, before 0 s, the β contour maps showed a power decrease in a wide area, especially for the precentral gyrus. After 0 s, the previous high γ increase disappeared and the β power increase was observed over the same wide area.

### 3.2. High γ and β band contour maps with mouth, water, and swallowing triggers

For each participant, an averaged normalized power contour map of the high γ and β band was created with mouth, water, and swallowing triggers (Fig.3). We could observe a different spatial pattern of power increase (white filled circles; the single-sided permutation test with Bonferroni correction, corrected *p* <0.05) in the high γ band within the three triggers. With the mouth trigger, a high γ burst was mainly observed in the precentral gyrus, and with the water trigger, the high γ burst was observed in the pericentral gyri, especially in the postcentral gyrus. For the swallowing, high γ bursts appeared in the region along the Sylvian fissure.

Conversely for the β band, it was difficult to identify a different spatial pattern. The power decrease (black filled circles; the single-sided permutation test with Bonferroni correction, corrected *p* <0.05) in the β band were widely observed for all three triggers.

### 3.3. Profiles of high γ and β band related to mouth, water, and swallowing triggers

#### 3.3.1. Power increasing vs. power decreasing

All contacts showing power increase or decrease in Figure 3 were plotted on the left hemisphere of the MNI normalized brain (Fig.4A). In the high γ band, the contacts of power increasing were plotted extensively in the water and swallowing group rather than the mouth group. In the β band, the contacts of power decreasing were plotted extensively across all groups.

In the high γ band, the numbers of contacts indicating power increasing were significantly larger than those indicating power decreasing in water and swallowing groups (Fig.4B) (the single-sided Wilcoxon signed-rank test with Bonferroni correction, corrected *p* = 0.012, and 0.023). In the β band, the numbers of contacts indicating power decreasing were significantly larger than those indicating power increasing in all three groups (the single-sided Wilcoxon signed-rank test with Bonferroni correction, *p* = 0.023, 0.012, and 0.0355) (Fig.4C).

We evaluated the differences across the three groups. The numbers of contacts indicating high γ burst were significantly different across the three groups (one-way ANOVA, *p* = 0.001) (Fig.4D). However, the normalized power of high γ burst (Fig.4E), the numbers of contacts indicating β attenuation (Fig.4F), and the normalized power of β attenuation (Fig.4G) showed no differences across groups (one-way ANOVA). These results indicated that high γ increasing and β decreasing were notably observed in all three groups, and different activities specific to mouth, water and swallowing triggers appeared in high γ increasing rather than β decreasing.

#### 3.3.2. Spatial distribution

The top 25% of contacts indicating a significant increase in high γ power or degree of β attenuation were plotted over the MNI brain (Fig.5). For the high γ power increasing, contacts related to water injection were active in the lateral portion of the central sulcus, and contacts related to swallowing were active in the regions along the Sylvian fissure. Contacts related to mouth opening were found in the precentral gyrus and in the ventrolateral prefrontal cortex (VLPFC), but were no better localized than the other groups.

In β power decreasing, contacts related to the mouth and water group were mainly localized in the VLPFC, and the localization appeared to be less specific in the swallowing group. This result indicated that high γ activities were better localized rather than β activities, and the high γ localization might reflect differences in neural processing.

#### 3.3.3. Dynamic power changes

For high γ and β band, we used three groups (Mouth-C, Water-C, and Swallow-C) and three trigger groups to derive nine different power plots (Fig.6). Data obtained from Mouth T, Water T, and Swallow T were arranged in chronological order.

**Fig 6.**
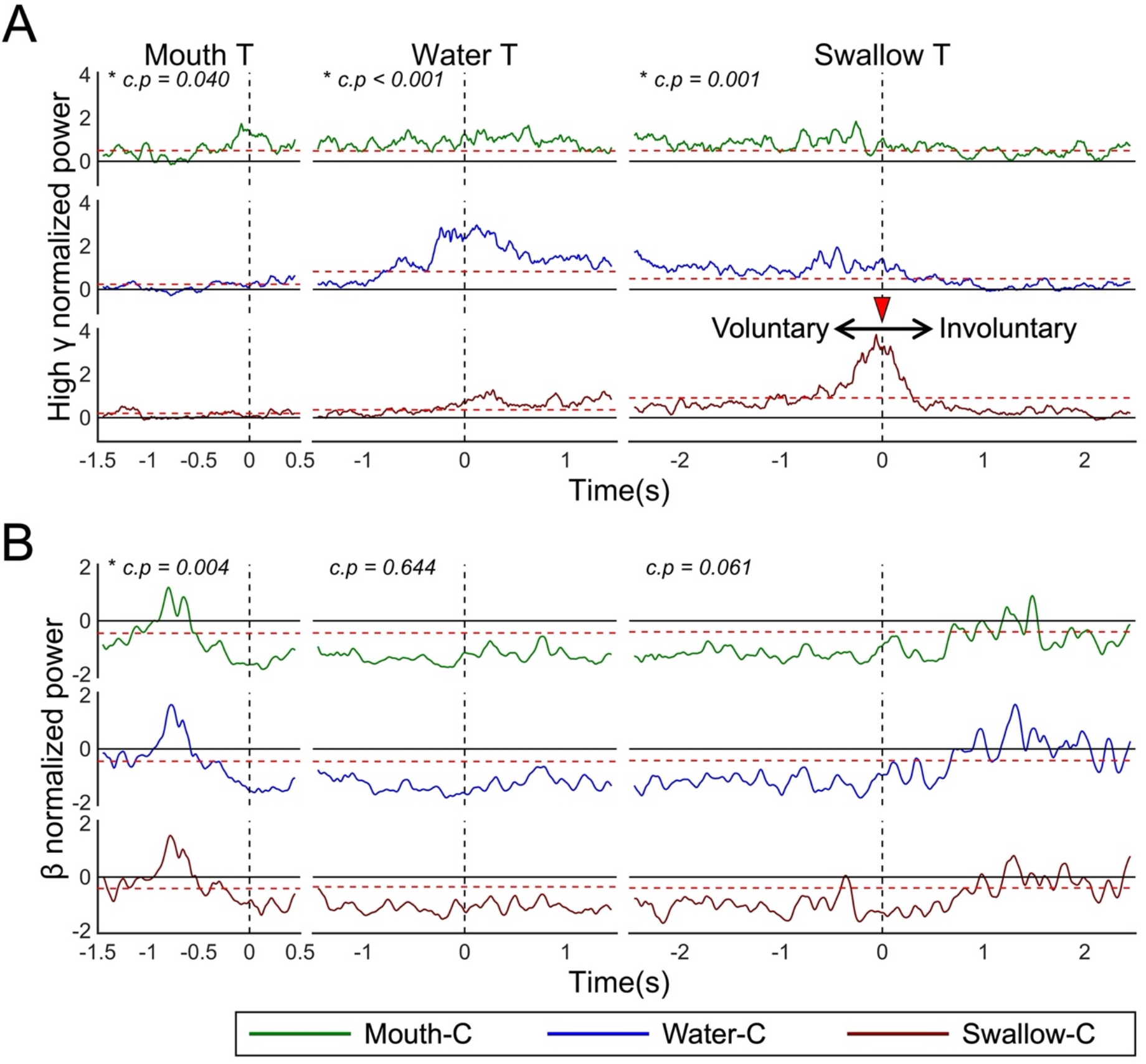
Temporal profile related to swallowing, mouth and water trigger groups. Averaged normalized power of high γ band (A) and β band (B) calculated from top 25% contacts, which showed significant power changes (Fig.5). The contacts group were indicated as Mouth-C, Water-C, and Swallow-C, colored by green, blue, and red in each. At 0 s of mouth triggers (Mouth T), only Mouth-C showed a high γ burst not Water-C and Swallow -C. At 0 s of Water T, Water-C showed notably high γ bursts. With Swallow T, Swallow-C showed high γ power increasing and achieved a peak at 0 s. After that, the power decreased. The 0 s of Swallow T corresponded to the transition time from voluntary swallowing to involuntary swallowing. In the β band, three contacts groups showed the same pattern as the β power decrease from Mouth T, and the attenuation was maintained until about 1.0 s of Swallow T. Red dot lines are FWE-corrected threshold. We evaluated the normalized power at 0 s of each trigger across Mouth-C, Water-C, and Swallow-C using one-way ANOVA. There were statistically significance in all triggers with high γ band and in Mouth T with β band.

Around 0 s of Mouth T, only Mouth-C showed a high γ burst. Around 0 s of Water T, Water-C showed a notable high γ burst. At 0 s of Swallow T, in Swallow-C, the high γ burst reached the peak value. In high γ activities, at 0 s of all triggers, there were significant differences across Mouth-C, Water-C, and Swallow-C (one-way ANOVA with Bonferroni correction, corrected *p* = 0.040 in Mouth T, corrected *p* < 0.001 in Water T, and corrected *p* = 0.001in Swallow T). The high γ burst showed statistically significance because they were bigger the FWE-corrected threshold (Fig.6A).

Moreover, the 0 s of the Swallow T corresponded to the boundary between voluntary swallowing and involuntary swallowing. Therefore, our results showed that during voluntary phase, high γ activities were increasing until completion of the voluntary swallowing, and subsequently, the high γ activities decreased.

Conversely, β power showed similar patterns for each trigger (Fig.6B). Except for Mouth T (one-way ANOVA with Bonferroni correction, corrected *p* = 0.004), there were no significant changes at 0 s of Water T and Swallow T across contacts groups (one-way ANOVA with Bonferroni correction). In Mouth T, β power was suppressed before 0 s. For Water T, β power remained inhibited, and with Swallowing T, β attenuation continued up to about 0.5 s, thereafter β power rebounded. The β attenuation showed statistically significance because they were lesser the FWE-corrected threshold.

#### 3.3.4. Time to change

We compared the lead-time when β or high γ power began to change between each trigger with each related contact. For the mouth trigger, β decreasing preceded the high γ increasing, but changes were not significant (the single-sided Wilcoxon signed-rank test) (Fig.7A). The lead-time of the high γ increasing showed significant differences across mouth, water, and swallowing group (one-way ANOVA, *p* = 0.004) (Fig.7B). The time of the swallow was earliest between three group. Even in the water, the beginning time of high γ burst preceded the 0 s of water triggers.

**Fig 7.**
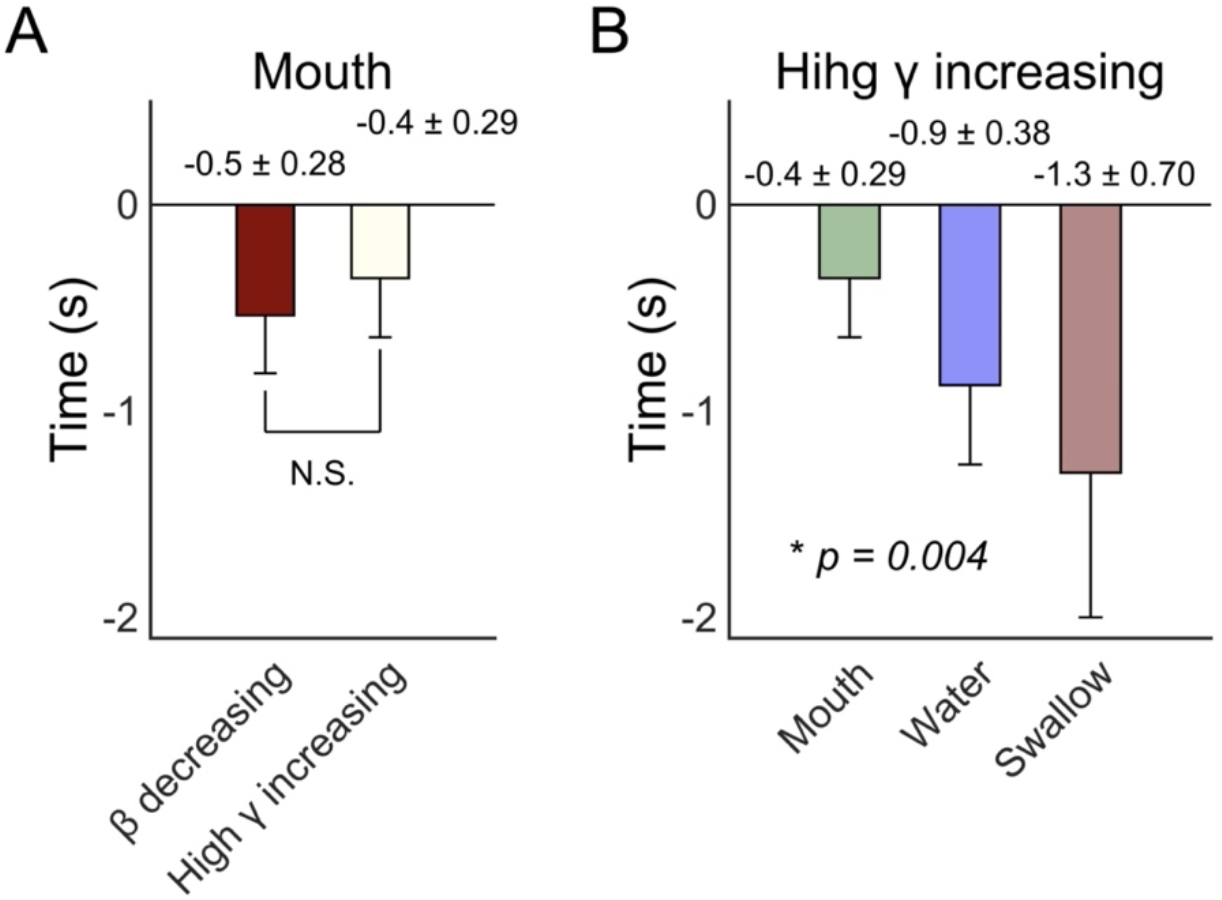
The beginning time of power changes. The beginning time of power changes for each band with each trigger was compared. In mouth triggers, there were no statistically differences between β decreasing and high γ increasing (single-sided Wilcoxon signed-rank test) (A). The time when swallowing-related high γ started to increase were earliest and statistically differences were observed across mouth, water, and swallowing triggers (one-way ANOVA) (B). The error bars indicate standard deviation.

### 3.4. β attenuation

To investigate β activities, we plotted the normalized β power from −1.5 s to 6 s of mouth triggers (Fig.8A). The β attenuation at 0 s of mouth triggers was maintained up to about 4 s, after which the β power rebounded. We investigated the correlation between β normalized power at 0 s of the mouth triggers (gray mesh area in Fig.8) – sequential β normalized power. The sequential correlation coefficients (r) were shown (Fig. 8B). Significant positive correlation above the threshold (red dashed line in Fig.8B) that meant 80 % statistical power were observed until rebound β appearance. The results indicated that the β attenuation was induced by mouth opening and the continuous β attenuation was correlated to the mouth-related β attenuation.

**Fig 8.**
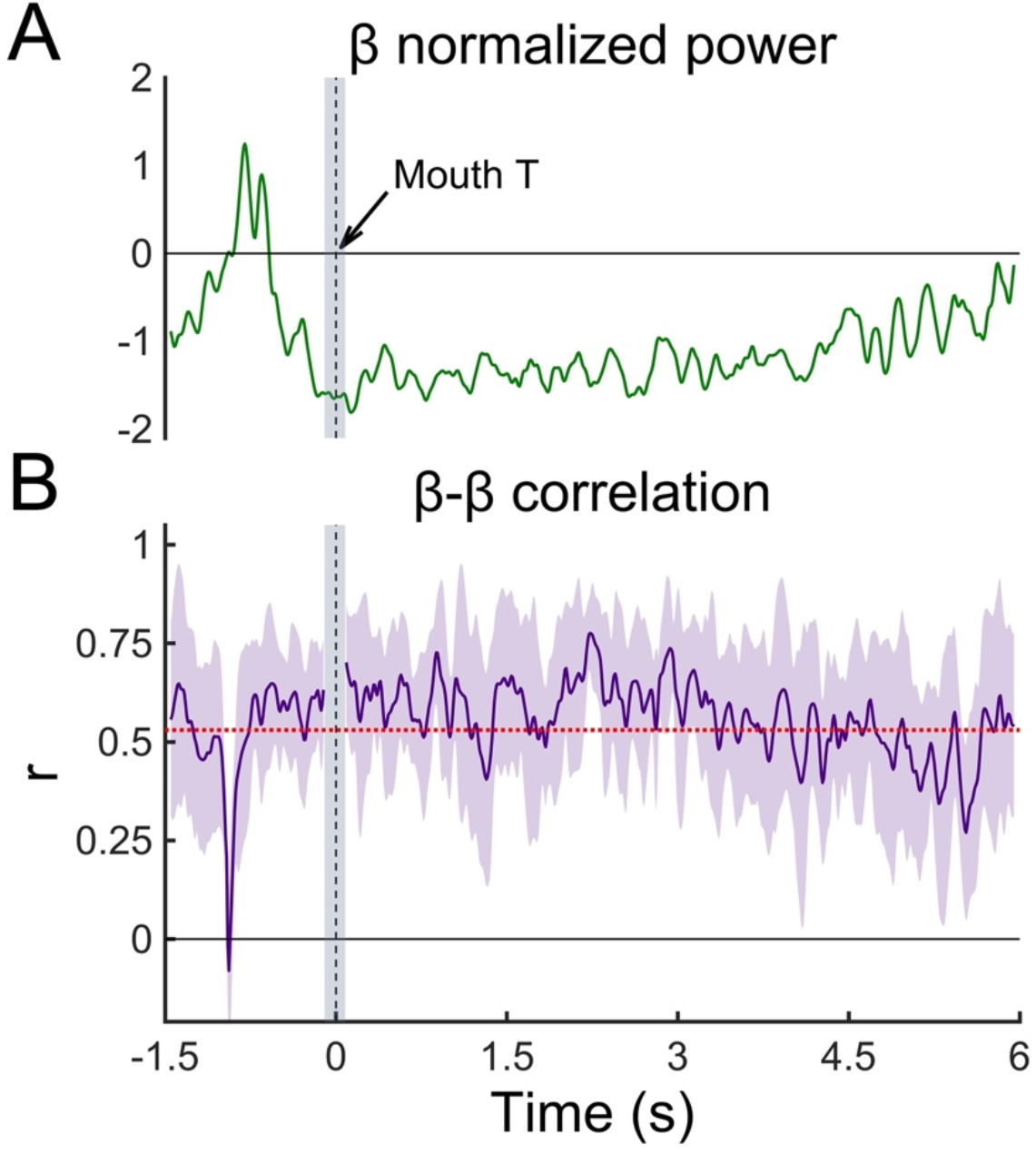
Mouth-related β band activities. Normalized power changes of β band with mouth triggers from −1.5 s to 6.0 s were presented (A). β normalized power was suppressed before 0 s, and remained inhibited, and, thereafter rebounded. Correlation coefficients (r) between β normalized power at 0 s (gray mesh area) and β normalized power at certain time during from −1.5 s to 6 s were shown (B). Red dashed line is the threshold that showed 80 % statistical power. During β attenuation, positive correlations that were above the threshold were observed. During β rebound, the positive correlations became weak. The error bars indicate 95% confidence intervals.

## 4 Discussion

This human ECoG study investigated orofacial cortical oscillatory changes up to the high γ band associated with swallowing. We demonstrated that different neural activities were observed between the high γ band and the β band. High γ burst and β attenuation were observed to be associated with mouth opening, water injection and swallowing, however, the distribution of high γ burst activity was more focal than that of β attenuation. Analyses based on mouth-related, water-related, and swallowing-related triggers showed statistically different high γ burst activity across these three groups. The key findings are that swallowing-related high γ activities appeared in the cortical areas along the Sylvian fissure and achieved the peak at the transition time from voluntary to involuntary swallowing.

Swallowing is divided into three phases, namely, the oral, pharyngeal, and esophageal phases. The oral phase is voluntarily controlled, whereas the pharyngeal and esophageal phases are involuntarily controlled (Jean, 2001). We inserted the swallowing trigger at the peak time of the EGG waveform, which corresponds to the boundary time between the oral and the pharyngeal phases (Kusuhara et al., 2004). Therefore, our swallowing triggers were inserted between the transition from voluntary to involuntary swallowing. In the present study, during the voluntary swallowing phase, the high γ bursts appeared; however, in conjunction with the completion of voluntary swallowing, the high γ activity also decreased. A previous ECoG study postulated that the cerebral cortex is involved in voluntary swallowing to a greater degree than in swallowing execution and that once the initial steps of volitional swallowing begin, the brainstem reflex is the primary drive for swallowing (Satow et al., 2004). Our finding that the high γ burst related to swallowing achieved a peak at the transition between voluntary and involuntary swallowing might indicate a neural mechanism: the main drive for swallowing switched from the cerebral cortex to the brainstem. Moreover, damage to cortical areas such as the Brodmann’s areas (BA) 43 and 44 are clinically known to cause dysphagia (Alberts et al., 1992; Baijens et al., 2008), and the opercular syndrome (Foix-Chavany-Marie syndrome) reveals that the voluntary phase of swallowing is severely affected, while reflex swallowing is preserved (Baijens et al., 2008). Taken together, these findings imply that the cortex plays a crucial role in execution of voluntary swallowing rather than involuntary swallowing.

High γ activities are involved in various neural activities, including somatosensory (Hirata et al., 2002), motor (Crone et al., 1998; Dalal et al., 2008), attention (Taylor et al., 2005), language (Hashimoto et al., 2017), and working memory (Pipa et al., 2009). High γ activities generally show more focal spatiotemporal distributions than lower frequency activities and more accurately reflect functional localization (Canolty et al., 2006; Crone et al., 1998; Dalal et al., 2008; Edwards et al., 2005). In the present study, we could demonstrate that using high γ activities, different cortical areas were activated associated with swallowing. Mouth opening-associated high γ burst appeared mainly in the lateral precentral gyrus and in the VLPFC, and these regions have been reported to be actively involved in orofacial motor action (Loh et al., 2020; Miller et al., 2007). Water-related high γ bursts were observed in the lateral portion along the central sulcus, which were known to be activated by tactile stimulation of the buccal mucosa (Miyaji et al., 2014). Swallowing-related high γ bursts were observed in regions along the Sylvian fissure, including the subcentral area (BA 43) and the frontal operculum (BA 44). Previous studies have reported that swallowing activated the subcentral area (Martin et al., 2004; Satow et al., 2004; Toogood et al., 2017a) and the frontal operculum (Dziewas et al., 2003; Hamdy et al., 1999a; Lowell et al., 2008).

Previous studies with fMRI (Hamdy et al., 1999a; Lowell et al., 2008; Martin et al., 2001; Mihai et al., 2014; Mihai et al., 2013; Toogood et al., 2017b) and PET (Hamdy et al., 1999b) have found that the caudolateral sensorimotor cortex (BA 1, 2, 3, and 4) and BA 43 and BA 44 are important in the preparation and execution of swallowing. However, we guessed that based on our results, the caudolateral sensorimotor cortex were activated by mouth-movement or sensory input from oral cavity and the areas activated by swallowing were BA 43 and BA 44. The caudolateral pre- and postcentral gyrus corresponds to the orofacial cortex (Martin et al., 1999; Salmelin et al., 1995), and tDCS and rTMS directly stimulate these cortical areas in order to induce neuronal excitation and promote beneficial neuroplasticity (Sasegbon et al., 2020). According to the present results, the cortical areas along the Sylvian fissure including BA 43 and BA 44 may have potential for novel targets of the neuromodulation that aims at recovery from dysphagia.

In our study, the subcentral area was activated by swallowing. However, the subcentral area was also activated by a communication function, such as vocal productions in response to acoustic perturbations (Chang et al., 2013) and eye-to-eye contact (Hirsch et al., 2017). The frontal operculum (BA 44) was also involved in swallowing, however, the BA 44 are is generally known as a key component of Broca’s language production region in humans (Loh et al., 2020). Monkeys possess an area comparable to human BA 44, and it has been hypothesized that BA 44 might have evolved as an area exercising high-level controls over orofacial activities including those related to communicative acts, and that, in the human brain, the BA 44 evolved to control the motor aspects of speech (Petrides et al., 2005). Therefore, it is feasible that both BA 43 and 44 are involved in communication.

In monkeys, the region anterior to the lateral sensorimotor cortex is known as the cortical masticatory area (CMA), and electrical stimulation of the CMA evoked swallowing (Martin et al., 1999). In the process of evolution, the early vertebrates already had acquired swallowing for feeding because the jawless vertebrates were able to swallow (Clark and Uyeno, 2019). Next, jawed vertebrates came to be able to masticate before swallowing. The orofacial muscular movements associated with mastication were eventually diverted to other functions, such as facial expressions for communication. Finally, in humans, orofacial muscular movements were used for motor language. Therefore, we think that the cortex associated with swallowing is quite extensive given the early acquisition in the evolutional process, and now overlaps areas that relate to mastication, facial expression, and speech which were originally involved in orofacial muscular movement.

The area of β attenuation was extensive, and β attenuation was similar for the mouth, water, and swallowing events. In a previous study, β attenuation in the lateral pericentral gyri was induced by the tongue movement and by swallowing (Dziewas et al., 2003). In our study, β attenuation was invoked associated with mouth opening, and the β band attenuation remained until the completion of the swallowing movement. Furthermore, continuous β band attenuation positively correlated with the mouth movement-associated β attenuation, therefore, we inferred that β attenuation observed in both of water and swallowing events were induced by mouth movements. Mouth-movement induced β attenuation simultaneously with high γ burst, which was consistent with the ECoG findings demonstrating that motor behaviour induces high γ activities along with decreases in β power (Crone et al., 1998; Dalal et al., 2008; Miller et al., 2007; Yanagisawa et al., 2012).

Moreover, this β attenuation was released after the completion of the swallowing movement and β power increased. This re-activation is known as rebounded β. MEG studies have reported that attenuation and subsequent rebound activities of 20 Hz were observed in the motor cortex and were attributed to mouth (Salmelin et al., 1995), lip and tongue movements (Salmelin and Sams, 2002). The β attenuation and subsequent rebound observed in this study agrees with these previous findings.

Somatosensory stimulation evokes high γ activities in the sensory motor cortex (Hirata et al., 2002), and we expected that water-associated high γ bursts would appear after the water triggers, however, contrary to our expectation, the beginning of the water-related high γ burst was 0.9 s earlier than the onset time of the water trigger. The onset time of the water trigger corresponded to the time when water bolus was injected by a syringe, which as fixed onto the lip. We inferred that the time lag was a result of the somatosensory input from the lip on which a syringe was placed. The high γ burst of the Water-C was observed notably only with Water T, and not at Mouth T and Swallow T. We concluded that the notably high γ bursts were evoked by somatosensory input from lip or mouth mucosa.

Our study has several limitations. First, we focused only the orofacial cortex, which is where the lateral region of the central sulcus is located. Multiple cortices are activated during swallowing (Ertekin and Aydogdu, 2003), however our study could not demonstrate cortical activities other than in orofacial region. Second, in this study, we divided swallowing events into three parts (mouth opening, water injection, and swallowing), and discussed motor and somatosensory neural processing. For more precise analysis, we may need to treat these as separate events, i.e., mouth movement only, water injection into the mouth only, and swallowing only. Third, we inferred the switching mechanism of the swallowing-related main driving force from the cortex to the brain stem. In the future, we think that simultaneous recording of the cortex and the brain stem may be needed to reveal how the swallowing-related interaction between them works.

## 5 Conclusion

With high γ activities, we distinguish the discrete cortical activities that are involved in swallowing and demonstrate that the cortical areas that are specific to swallowing are the regions along the Sylvian fissure including the subcentral area and the frontal operculum. Swallowing-related high γ activities that achieved the peak at the boundary time between voluntary and involuntary swallowing may represent the neural processing of switching from the cortex to the brainstem involved in swallowing execution.

## Abbreviations

CPG: central pattern generator
rTMS: repetitive transcranial magnetic stimulation
tDCS: transcranial direct current stimulation
BMI: brain-machine interface
EEG: electroencephalogram
PET: positron emission tomography
NIRS: near-infrared spectroscopy
fMRI: functional magnetic resonance imaging
MEG: magnetoencephalography
ECoG: electrocorticogram
CT: computed tomography
MNI: Montreal Neurological Institute
EGG: electroglottograph
Mouth-C: mouth-related contacts
Water-C: water-related contacts
Swallow-C: swallowing-related contacts
Mouth T: mouth triggers
Water T: water triggers
Swallow T: swallowing triggers
FWE: family-wise error
ANOVA: analysis of variance
Ch: channel
VLPFC: ventrolateral prefrontal cortex
BA: Brodmann’s areas
CMA: cortical masticatory area.

## Author Contributions

M.H. designed the study. H.H. performed the experiments, assisted with the epileptic medical treatment, created the MATLAB program and analyzed the data, created all the figures and tables, and was primarily responsible for writing the manuscript. F.Y. H.M. and M.H. assisted in the acquisition of the measurements. S.K. developed some devices to help with the measurements. H.K., S.O., N.T., H.M.K. and M.H. performed the epilepsy surgery, and Ta.Ya. assisted with the epileptic medical treatment. M.H., K.T. and Ta.Ya advised H.H. on scientific matters. M.H. and K.T. revised the manuscript. To.Yo., H.K. and M.H. supervised the experiments and analyses. All authors reviewed the manuscript.

